# MYB12 spatiotemporally represses TMO5/LHW-mediated transcription in the Arabidopsis root meristem

**DOI:** 10.1101/2022.03.08.483486

**Authors:** Brecht Wybouw, Helena E. Arents, Baojun Yang, Jonah Nolf, Wouter Smet, Michael Vandorpe, Daniël Van Damme, Matouš Glanc, Bert De Rybel

**Affiliations:** Ghent University, Department of Plant Biotechnology and Bioinformatics, Technologiepark 71, 9052 Ghent, Belgium; VIB Centre for Plant Systems Biology, Technologiepark 71, 9052 Ghent, Belgium

**Author notes:** Corresponding author. (B.D.R.). These authors contributed equally (Brecht Wybouw, Helena E. Arents). Co-senior authors (Matouš Glanc, Bert De Rybel).

## Abstract

Transcriptional networks are crucial to integrate various internal and external signals into optimal responses during plant growth and development. Primary root vasculature patterning and proliferation are controlled by a network centred around the basic Helix-Loop-Helix transcription factor complex formed by TARGET OF MONOPTEROS 5 (TMO5) and LONESOME HIGHWAY (LHW), which control cell proliferation and orientation by modulating cytokinin response and other downstream factors. Despite recent progress, many aspects of the TMO5/LHW pathway are not fully understood. In particular, the upstream regulators of TMO5/LHW activity remain unknown. Here, using a forward genetic approach to identify new factors of the TMO5/LHW pathway, we discovered a novel function of the MYB-type transcription factor MYB12. MYB12 physically interacts with TMO5 and dampens the TMO5/LHW-mediated induction of direct target gene expression as well as the periclinal/radial cell divisions. The expression of *MYB12* is activated by the cytokinin response, downstream of TMO5/LHW, resulting in a novel MYB12-mediated negative feedback loop that restricts TMO5/LHW activity to ensure optimal cell proliferation rates during root vascular development.

## Introduction

Transcription factors (TFs) play a crucial role in controlling virtually all developmental processes in eukaryotes by regulating the expression of specific subsets of target genes. TFs do not typically act alone but are embedded in complex transcriptional networks, which modulate their activity to ensure optimal transcriptional output in response to various environmental and developmental signals. Transcriptional networks often rely on feedback regulation, where a TF promotes the expression of its own activator (positive feedback) or of its repressor (negative feedback), respectively (Ohashi-Ito and Fukuda, 2020).

During vascular development in the plant embryo and primary root apical meristem, the heterodimer complex formed by the basic Helix-Loop-Helix (bHLH) TFs TARGET OF MONOPTEROS 5 (TMO5) and LONESOME HIGHWAY (LHW) controls vascular cell proliferation leading to radial expansion of the vascular bundle (Ohashi-Ito and Bergmann, 2007; De Rybel et al., 2013; Ohashi-Ito et al., 2013; De Rybel et al., 2014; Ohashi-Ito et al., 2014). The TMO5/LHW dimer is active in xylem cells, where it directly activates the expression of *LONELY GUY 3* (*LOG3*), *LOG4* and *BETA GLUCOSIDASE 44* (*BGLU44*), encoding key enzymes in the biosynthesis and deconjugation of cytokinin (Kurakawa et al., 2007; Kuroha et al., 2009; De Rybel et al., 2014; Ohashi-Ito et al., 2014; Yang et al., 2021). This leads to a local increase of cytokinin, which is thought to diffuse to the neighbouring procambium cells (De Rybel et al., 2014; Ohashi-Ito et al., 2014) and trigger the expression of members of the DNA-BINDING WITH ONE FINGER (DOF) type TF family (Miyashima et al., 2019; Smet et al., 2019). These DOF-type TFs in turn lead to a switch in division plane orientation from anticlinal to periclinal and radial in specific subsets of procambium and phloem pole cells, depending on the DOF family member. The actual molecular mechanisms are however not yet fully explored (Otero S., 2021). The activity of the TMO5/LHW complex is negatively regulated by members of the SUPPRESSOR OF ACAULIS51-LIKE (SACL) subclade of bHLH TFs (Katayama et al., 2015; Vera-Sirera et al., 2015). Similarly, to TMO5, SACLs physically interact with LHW. By competing with TMO5 for LHW binding, the SACLs reduce the amount of functional TMO5/LHW complexes, and thus dampen the activity of the pathway (Katayama et al., 2015; Vera-Sirera et al., 2015). As *SACL* genes are themselves downstream targets of TMO5/LHW, they constitute a typical negative feedback loop (Katayama et al., 2015; Vera-Sirera et al., 2015).

Besides forming bHLH homo-or heterodimers, bHLH proteins have also been shown to directly interact with other proteins such as MYB-type TFs, which can enhance or supress their transcriptional activity (Zhao et al., 2008; Carretero-Paulet et al., 2010; Feller et al., 2011; Cui et al., 2021). MYB TFs are defined by their highly conserved DNA-binding MYB-domain that contains up to four α-helical “R” repeats (Ogata et al., 1996; Du et al., 2009). The class (R1, R2 or R3, depending on their similarity to c-Myb R repeats) and number of the R repeats are the basis of MYB protein classification (Dubos et al., 2010). Most plant MYBs belong to the R2R3-MYB subfamily (Stracke et al., 2001), which is involved in a plethora of processes including phenylpropanoid biosynthesis (Liu et al., 2015), development of tissues and organs (Oppenheimer et al., 1991; Lee and Schiefelbein, 1999) and hormonal responses (Jin and Martin, 1999). Exemplary bHLH-MYB interactions take place during epidermal cell fate specification. The formation of trichomes and root hairs depends on the assembly of different heterotrimeric bHLH/WD40/MYB complexes. In addition to the WD40 protein TRANSPARENT TESTA GLABRA 1 (TTG1), the core bHLH proteins GLABRA 3 (GL3) or ENHANCER OF GLABRA 3 (EGL3) interact with the R2R3 MYB proteins WEREWOLF (WER) or GLABRA 1 (GL1), forming an active transcriptional complex that promotes root hair or trichome formation, respectively. Alternatively, the recruitment of CAPRICE (CPC), TRIPTYCHON (TRY) or ENHANCER OF TRY AND CPC 1, single-repeat R3 MYBs that lack the C-terminal transcriptional activation domain and compete with the transcriptional activating R2R3 MYBs for bHLH binding, results in the formation of a transcriptional inactive complex that prevents trichome/root hair formation (Wada et al., 1997; Kirik et al., 2004; Ramsay and Glover, 2005; Tominaga-Wada et al., 2017). The single-repeat R3 MYBs are downstream targets of the active MYB/bHLH/WD40 complex, and at the same time its non-cell autonomous inhibitors. The bHLH and MYB TFs thus constitute a negative feedback loop that lies at the core of epidermal cell type specification and patterning (Wang et al., 2008; Song et al., 2015). A similar bHLH/MYB/WD40 complex controls the expression of a core enzyme in the proanthocyanin biosynthetic pathway (Appelhagen et al., 2011; Xu et al., 2013; Xu et al., 2015). As such, interactions between MYB and bHLH TFs are key to various developmental processes.

The closely related R2R3 MYB proteins MYB11, MYB12 and MYB111 promote the expression of genes encoding key flavonol biosynthetic enzymes (Mehrtens et al., 2005; Stracke et al., 2007; Stracke et al., 2010; Stracke et al., 2017). Flavonols are a subgroup of flavonoids, besides the red to purple anthocyanins and brown proanthocyanidins (Winkel-Shirley, 2001; Lepiniec et al., 2006). Flavonoids convey color to fruits and seeds and aid in abiotic stress response (Wang et al., 2016). MYB11, MYB12 and MYB111 induce flavonol biosynthesis at different developmental stages, depending on their distinct expression patterns: While MYB12 is mostly active in roots, MYB11 acts in meristematic tissues and MYB111 functions in the hypocotyl and cotyledons (Stracke et al., 2007). The genes encoding flavonol biosynthesis enzymes *CHALCONE SYNTHASE* (*CHS*), *CHALCONE FLAVANONE ISOMERASE* (*CHI*), *FLAVANONE 3’-HYDROXYLASE* (*F3’H*), and *FLAVONOL SYNTHASE* (*FLS*) catalyse consecutive steps of flavonol production (Forkmann and Martens, 2001) and are regulated by MYB TFs via the MYB recognition element in their promoter regions. *CHS* and *FLS* are directly transcriptionally activated by MYB12 (Mehrtens et al., 2005). Consequently, the levels of the flavonols kaempferol and quercetin are decreased in the *myb12* mutant, while *MYB12* overexpression leads to increased flavonol levels (Mehrtens et al., 2005).

Here, we discover a novel role of MYB12 as a negative regulator of the TMO5/LHW pathway during vascular proliferation. *MYB12* is a downstream target of *TMO5/LHW*; interacts with TMO5 and represses TMO5/LHW transcriptional activity, thus constituting a negative feedback loop in the regulation of vascular development. Our work highlights the importance of bHLH-MYB interactions in multiple developmental processes; and demonstrates concomitant activator and repressor functions of the same TF in different transcriptional network contexts.

## Results

### A mutant screen identifies modulators of TMO5/LHW activity

In order to identify novel regulators of TMO5/LHW activity leading to vascular proliferation via control of oriented cell divisions, we designed an EMS-based forward genetic screen in the previously described dexamethasone (DEX)-inducible p*RPS5A*::TMO5:GR x p*RPS5A*::LHW:GR double misexpression line (double GR or dGR) in Col-0 background (Smet et al., 2019). Upon exogenous DEX treatment, root apical meristem width is increased in this line due to the ectopic periclinal and radial cell divisions (De Rybel et al., 2013), protoxylem differentiation is inhibited due to increased cytokinin levels (De Rybel et al., 2014) and additionally, primary root length is reduced (De Rybel et al., 2013) (**Fig. 1A-H, Table S1**). We reasoned those mutations in positive/negative regulators of the TMO5/LHW pathway would suppress/enhance these dGR phenotypes. Although the TMO5/LHW activity was previously shown by a detailed quantification of the vascular cell file number (Arents et al., 2022), such experiments are labour intensive and require fixed samples, making them incompatible with high-throughput screening. We thus first evaluated whether root length and meristem width could serve as reliable read-outs for TMO5/LHW activity and hence vascular cell proliferation capacity, by plotting the root length or root width parameters against the number of quantified vascular cell files in multiple transgenic lines with increasing levels of TMO5/LHW heterodimer activity (Col-0, p*RPS5A*::LHW:GR, p*RPS5A*::TMO5:GR, the inducible dGR line and a constitutive double TMO5/LHW misexpression line). We observed a clear inverse correlation between root length and TMO5/LHW activity and a positive correlation between root width and TMO5/LHW activity (**Fig. 1I-J, Table S1**). These results suggest that root length and width can serve as reliable proxies for the number of vascular cell file number and thus TMO5/LHW activity.

**Figure 1.**
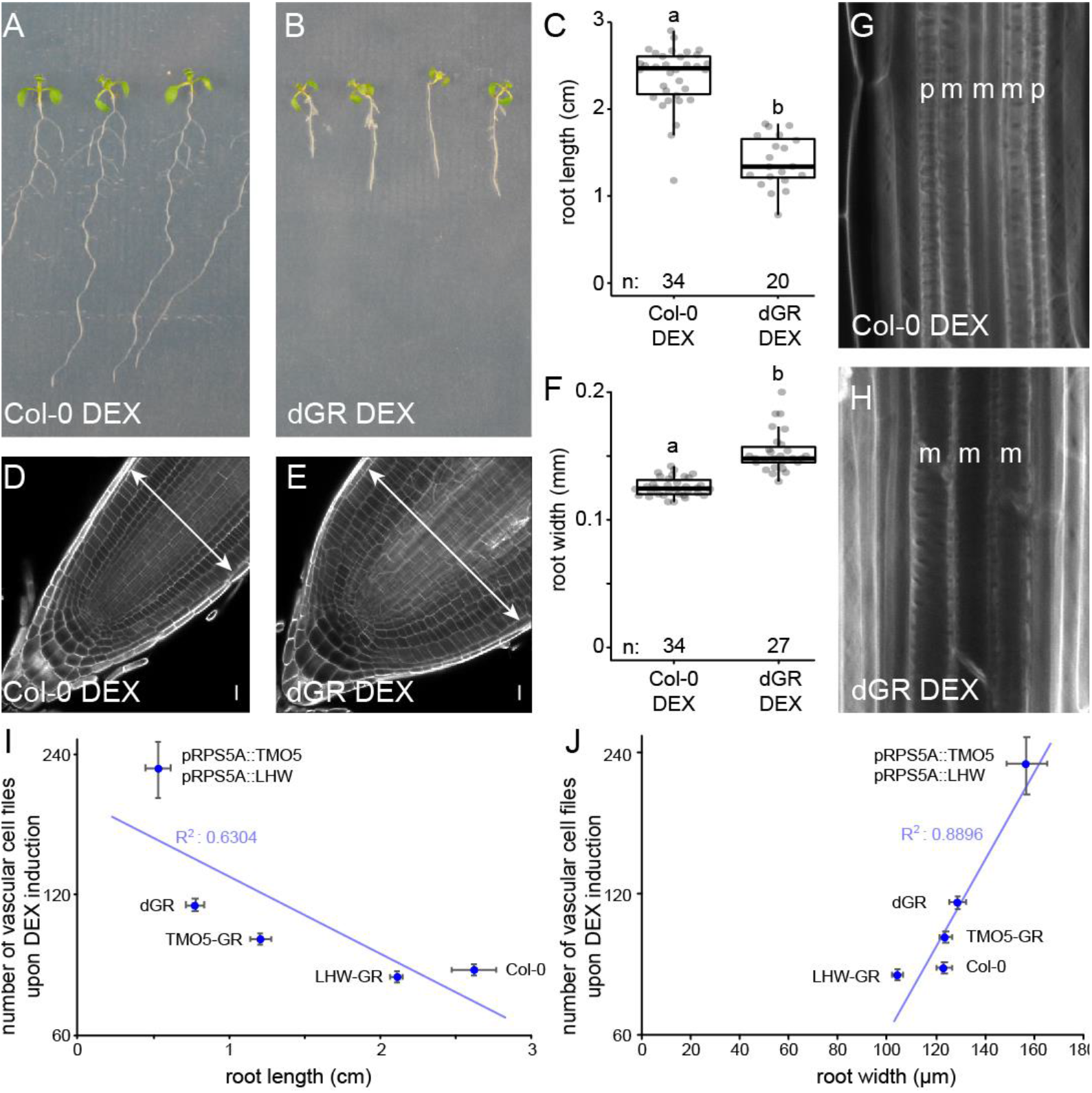
Root phenotype of Col-0 and the dGR line on induced (DEX) media. (**A**-**B**) 1-week-old Col-0 (**A**) and dGR (**B)** plants grown on 10 µM DEX. (**C**) Boxplot of root length of Col-0 and dGR plants grown for 5 days on 10 µM DEX. (**D**-**E**) Col-0 (**D**) and dGR (**E**) root tips grown on 10 µM DEX. Arrows are highlighting root meristem width. (**F**) Boxplot of root width of Col-0 and dGR plants grown on 10 µM DEX. (**G-H**) The vascular differentiation phenotype of Col-0 (**G**) and dGR (**H**) plants grown on 10 µM DEX. The p and m indicate protoxylem and metaxylem strands respectively. Root width of Col-0 and dGR plants grown for 5 days on 10 µM DEX (n ≥ 20). (**I**-**J**) 1-week old seedlings grown on 10 µM DEX (n ≥ 10), were used to plot the number of total cell files in the root meristem against the root length (**I**) or root width (**J**). Error bars indicate standard error. Scale bars in D-E indicate 10 µm. Lower-case letters in C, F indicate significantly different groups as determined by one-way ANOVA with post-hoc Tukey HSD testing. Black lines indicates mean values and grey boxes indicate data ranges. n marks the number of datapoints for each sample.

Having established the screening strategy, we performed EMS mutagenesis of dGR seeds and screened 228 pools of EMS mutagenized M_2_ dGR seedlings for alterations in root length upon DEX induction. This first round of selection yielded 310 candidate mutants from 110 pools, of which 260 produced viable M_3_ seeds. In total, 50 albino plants were observed among these 228 pools of mutants, suggesting that the EMS mutagenesis was successful (Micol-Ponce et al., 2014). In the M_3_ generation, we quantified both root length and root meristem width of the 260 candidate mutants (**Fig. 2**), resulting in 20 validated mutants with reduced responses (*insensitive 1-20, ins1-20*) and 2 mutants showing hypersensitive responses (*hypersensitive 1-2, hyp1-2*) (**Fig. S1-3, Table S1**). We next performed a detailed quantification of the vascular cell file number as the read-out of TMO5/LHW activity used previously (Ohashi-Ito and Bergmann, 2007; De Rybel et al., 2013; Ohashi-Ito et al., 2013; De Rybel et al., 2014; Ohashi-Ito et al., 2014). Notably, 8 insensitive and 1 hypersensitive mutants already showed a respectively significantly reduced number of vascular cell files in mock conditions (**Fig. 3, Table S1**), suggesting that these mutants might inherently have differential TMO5/LHW activity and further confirming that our multi-step screening procedure using root length and width as proxies was successful. A segregation analysis further showed that the observed phenotypes in *ins2* and *ins7* could not be explained by a recessive mutation at a single locus (**Table S2**). These mutants were therefore excluded from further analysis. We finally focussed our attention to the mutants with the most pronounced phenotype in each category: *ins4* and *hyp2* (**Fig. 3, Table S1**), and mapped the causal mutations by next generation sequencing.

**Figure 2.**
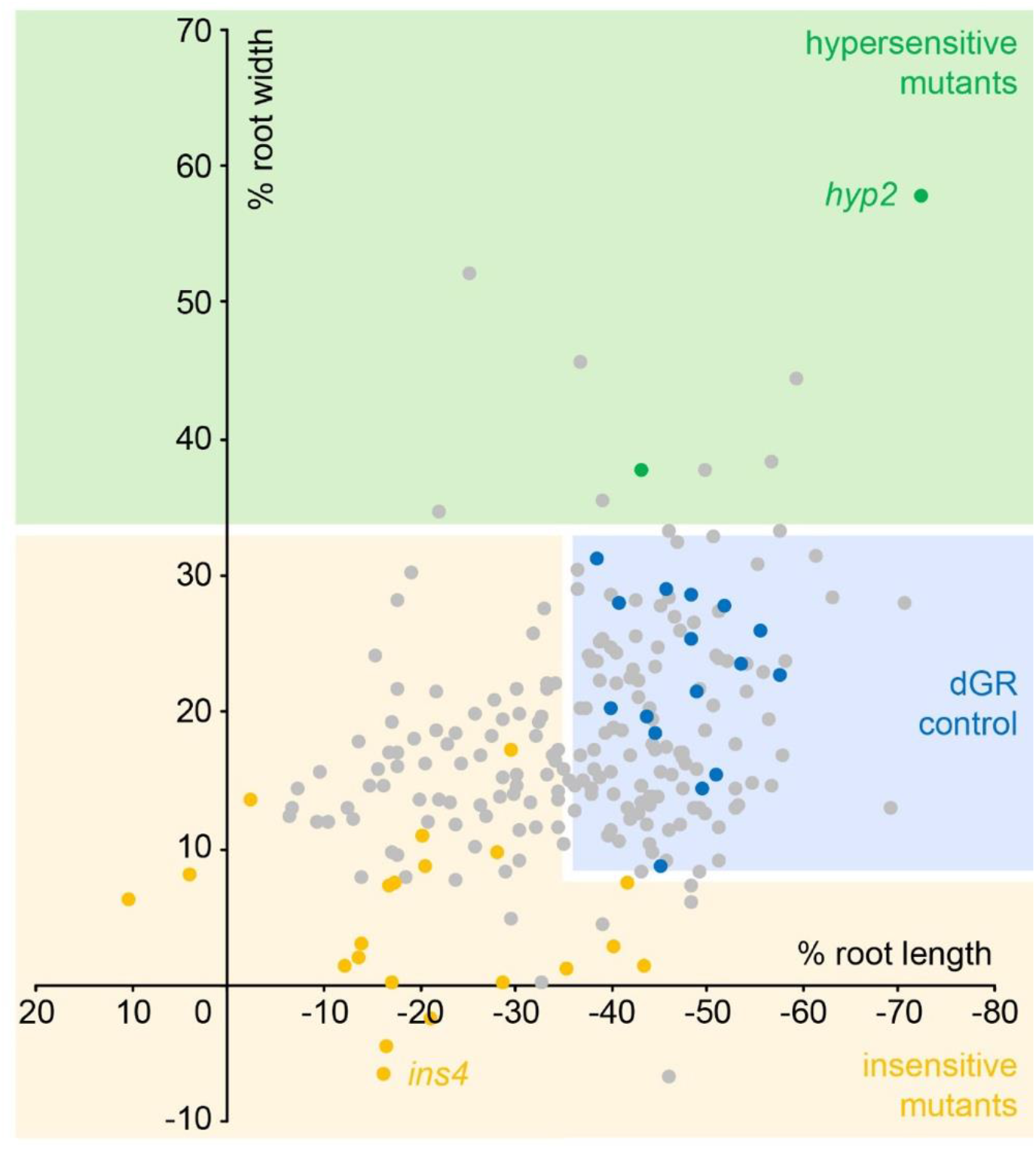
Overview of obtained EMS mutants. A total overview of all 260 primary selected EMS mutants is plotted for their sensitivity of root length changes relative to dGR against the sensitivity of root width changes relative to dGR. Data from the EMS screening was used. Dots in the blue box represent EMS mutants behaving similar to parental dGR control and dots in the yellow box represent mutants that behave insensitive to dGR response compared to the dGR parental line, while in the green box mutants behave hypersensitive to dGR induction. Yellow and green dots represent the 22 selected EMS mutants, the yellow dots represent the *ins* mutants and green dots the *hyp* mutants. Grey dots represent other EMS mutants selected from the primary screen and blue dots represent non-mutagenized parental dGR. For each data point the average was used from 10 biological repeats.

**Figure 3.**
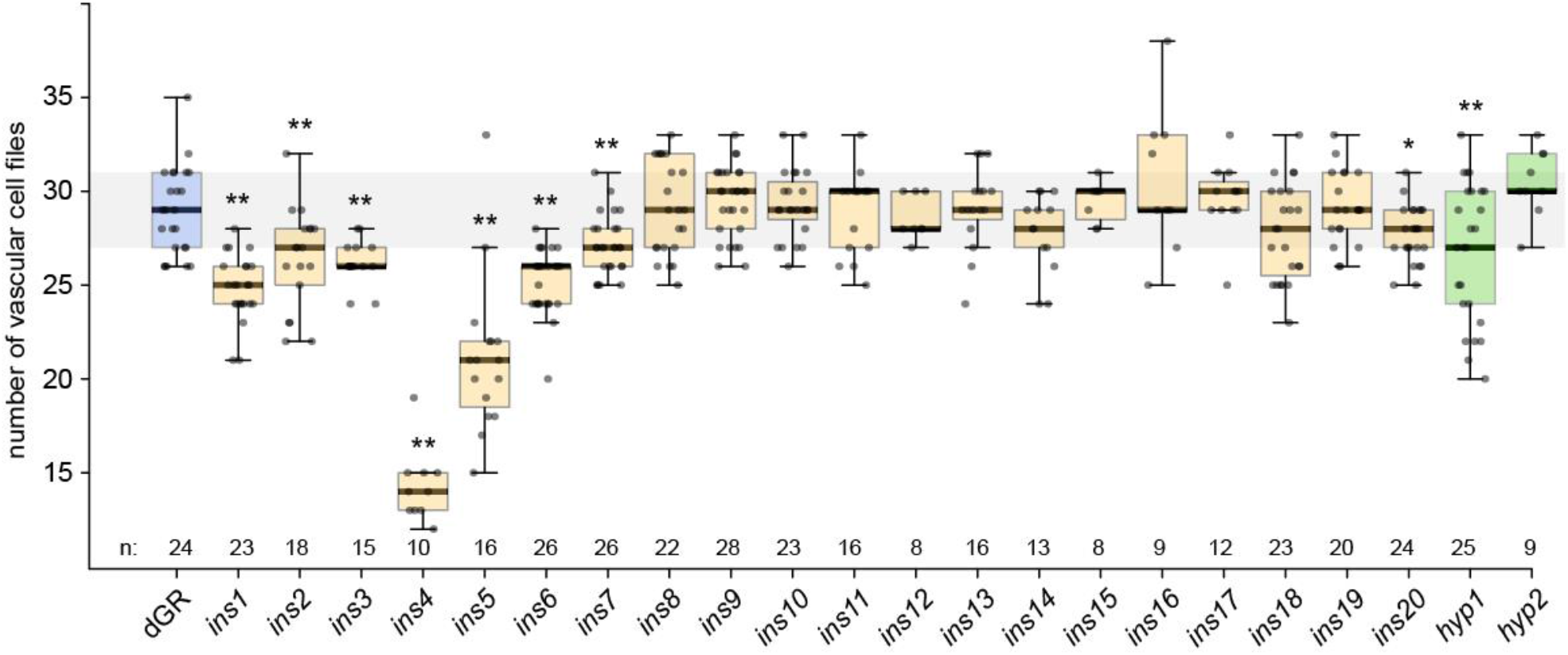
Overview vascular cell files phenotype in candidate mutants. Counts of vascular cell files in the root meristem of 1-week-old dGR (blue), *ins* (yellow) and *hyp* (green) seedlings. n marks the number of datapoints for each sample. Student’s T-test was performed to evaluate statistical differences between a mutant’s and dGR vascular cell file numbers. Student T-test significances asterisks: * = p-value < 0.05; ** = p-value < 0.01.

### A strong lhw allele is causal to the ins4 phenotype

The insensitive *ins4* mutant showed a strong reduction in the number of vascular cell files under mock condition and an almost complete repression of the increased root thickness upon DEX treatment (**Fig. 3, Fig. S1, Table S1**). Sequencing and SHORE map analysis (Schneeberger et al., 2009) revealed that *ins4* carried a premature stop codon in *LHW* (**Fig. 4A**). Similar to published *lhw* mutant alleles (Ohashi-Ito and Bergmann, 2007; Parizot et al., 2008; De Rybel et al., 2013; Ohashi-Ito et al., 2013), the *ins4* mutant showed a monarch vascular architecture in the primary root meristem, resulting in an off-centre xylem bundle during secondary growth (**Fig. 4B-H, Table S1**). The number of vascular cell files could also be rescued by exogenous cytokinin application (**Fig. 4I, Table S1**) as was shown before to bypass the TMO5/LHW dependent cytokinin biosynthesis (De Rybel et al., 2014). Taken together, the mapping and phenotypic characterization show that *ins4* is a novel, strong *lhw* allele. We thus termed *ins4* as *lhw-8*. As TMO5/LHW activity is highly dose-dependent (De Rybel et al., 2013; Ohashi-Ito et al., 2013; De Rybel et al., 2014; Ohashi-Ito et al., 2014; Smet et al., 2019), mutations in TMO5 or LHW were an expected outcome of our screen. Therefore, although *ins4 / lhw-8* itself does not provide new insight into the regulation of TMO5/LHW activity, it further confirms that our screening set-up was successful and yielded relevant mutants.

**Figure 4.**
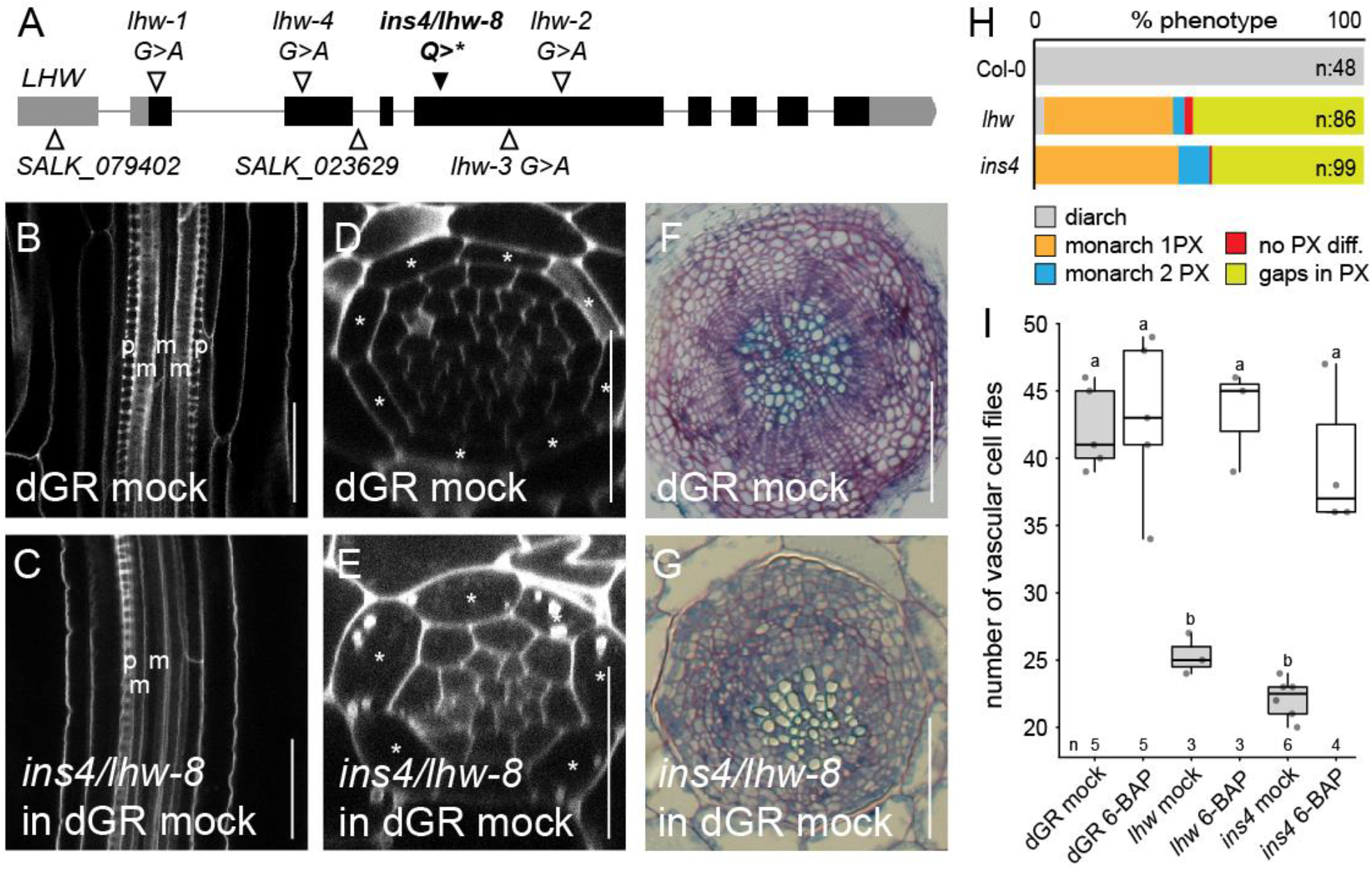
The insensitive mutant *ins4* is a novel *lhw* allele. (**A**) Alleles of *lhw* mutants with *ins4*/*lhw-8* having a point mutation, resulting in a premature stop codon in exon (black bar) 4 of *LHW*. (**B**-**C**) Longitudinal view of the root vascular tissue shown for Col-0 (**B**) and *ins4*/*lhw-8* (**C**). (**D-E**) Optical cross-section through the root meristem of Col-0(**D**) and *ins4*/*lhw-8* (**E**) show smaller vascular cylinder for *ins4*/*lhw-8*. (**F**-**G**) Secondary growth phenotype can be observed in sections of Col-0 (**F**) and *ins4/lhw-8* (**G**) through the hypocotyl of 3-week-old seedlings. Scale bars in B-E are 25 µm and in F-G 100 µm. (**H**) The frequency of xylem differentiation (diff.) phenotype plotted for Col-0, *lhw* and *ins4*. The asterisks mark the endodermis cells in D-E, ‘p’ an ‘m’ represent protoxylem and metaxylem cell files in B-C. (**I**) The number of vascular cell files of 1-week-old seedlings treated with cytokinin (6-BAP). Lower-case letters indicate significantly different groups as determined by pairwise comparison in a two-way ANOVA. Black lines indicates mean values and grey/white boxes indicate data ranges. n marks the number of datapoints for each sample.

### hyp2 is a novel myb12 allele

At the other side of the selected mutant spectrum, the recessive *hyp2* mutant showed little or no aberrant phenotype under normal growth conditions, but a strong hypersensitive response upon DEX treatment (**Fig. 3, Fig. 5N, Fig. S1, Table S1**). SHORE map analysis (Schneeberger et al., 2009) identified an early stop codon in the gene encoding the R2R3 transcription factor MYB12 (**Fig. 5A**). To confirm the causality of the *MYB12* mutation for the observed dGR hypersensitive phenotype, we first crossed the previously published *myb11 myb12-1f myb111* triple mutant (Stracke et al., 2007) into our dGR parental line. A hypersensitive response comparable to *hyp2* was detected in the *dGR myb11 myb12-1f myb111* mutant (**Fig. 5B-I, N, Table S1**). This triple mutant also did not show an aberrant phenotype under mock conditions in the Col-0 control background (**Fig. 5B, H, N, Table S1**). Next, we complemented the *hyp2* mutant with a construct driving the *MYB12* coding sequence from the meristematic *RPS5A* promoter (Weijers et al., 2001). The p*RPS5A*::MYB12 line showed a mild repression of the number of vascular cell files in mock conditions, which was correlated with *MYB12* expression levels as determined by qRT-PCR analysis (**Fig. S4A-B, Table S1**). Upon DEX treatment, p*RPS5A*::MYB12 construct strongly repressed the TMO5/LHW induced vascular cell proliferation (**Fig. 5J-N, Table S1**). Taken together, *hyp2* is a novel mutant allele of *MYB12*, which we designated as *myb12-2*. Our initial results hint towards a new function for this TF and suggest that MYB12 might act as a negative regulator of the TMO5/LHW pathway.

**Figure 5.**
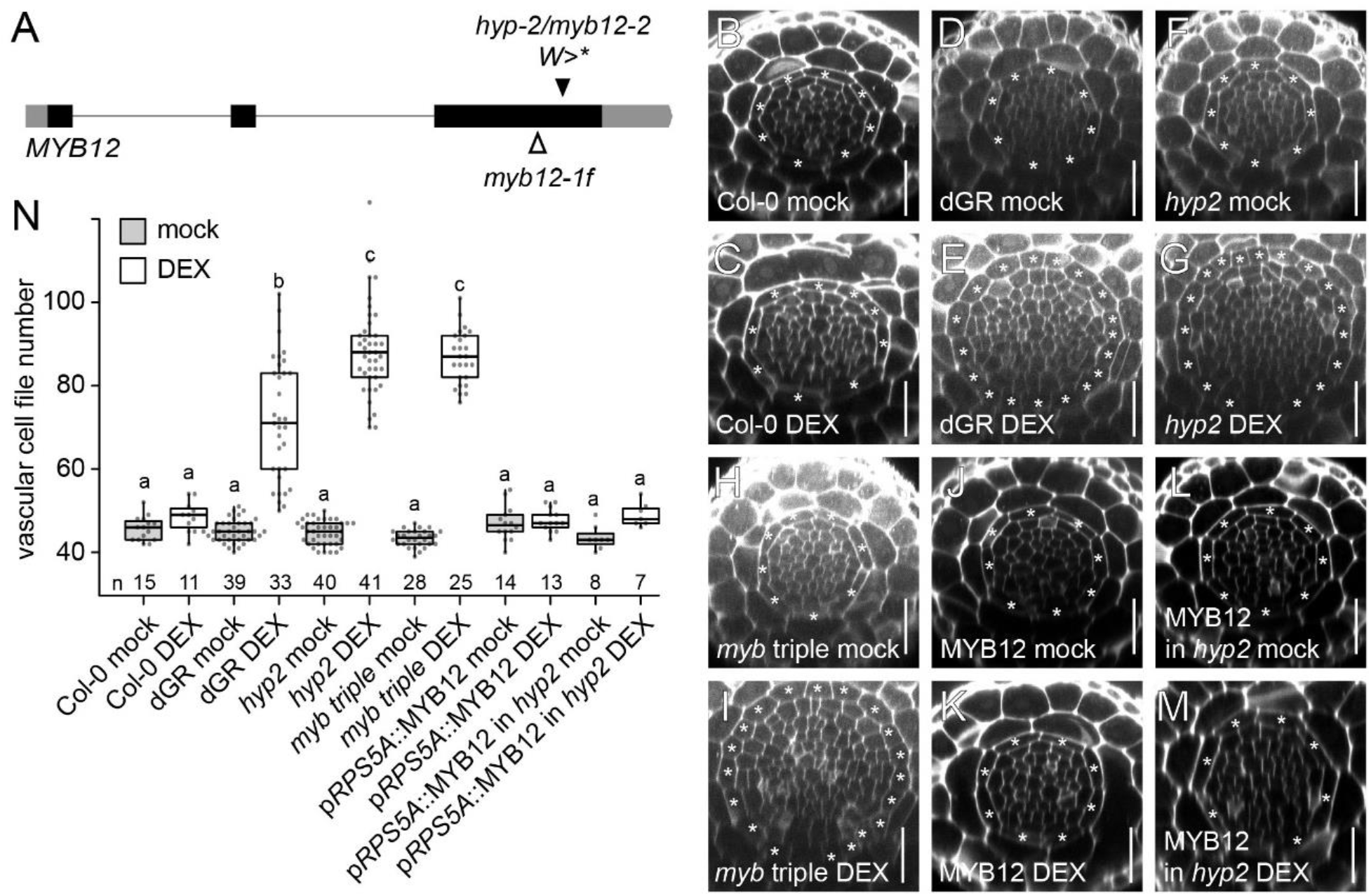
The *hyp2* is hypersensitive to dGR response and MYB12 acts as a repressor for TMO5/LHW activity. (**A**) MYB12 gene marked with known *myb12-1f* transposon insertion site and hyp2/*myb12-2* point mutation site, which results in premature stop codon. (**B**-**M**) Representative root meristem cross-sections of Col-0 on mock (**B**), Col-0 on DEX (**C**), dGR on mock (**D**), dGR on DEX (**E**), *hyp2***/**myb12-2 on mock (**F**), *hyp2***/**myb12-2 on DEX (**G**), *myb11 myb12-1f myb111* triple mutant (referred to as *myb triple*) on mock (**H**), myb triple on DEX (**I**), p*RPS5A*::MYB12 on mock (**J**), p*RPS5A*::MYB12 on DEX (**K**), p*RPS5A*::MYB12 (in *myb12-2*) line on mock (**L**) and on DEX (**M**). The asterisks mark the endodermis cells and counted vascular cell file number are within this cell type. Scale bars are 25 µm. (**N**) Boxplot plotting the vascular cell file number. Lower-case letters indicate significantly different groups as determined by pairwise comparison in a two-way ANOVA. Black lines indicates mean values and grey/white boxes indicate data ranges. n marks the number of datapoints for each sample.

Additionally, we previously found that *MYB12* is transcriptionally upregulated upon TMO5/LHW induction in the dGR line (Smet et al., 2019) (**Fig. 6A**) and validated this result by qRT-PCR analysis (**Table S1)**. Given the slow induction kinetics compared to direct TMO5/LHW target genes such as *LOG4* (De Rybel et al., 2014; Ohashi-Ito et al., 2014), we hypothesized that the induction of *MYB12* is likely indirect and possibly triggered by cytokinin signalling downstream of TMO5/LHW (De Rybel et al., 2014; Ohashi-Ito et al., 2014). Indeed, we found *MYB12* to be cytokinin inducible by qRT-PCR analysis (**Fig. 6B, Table S1**), confirming previous reports (Brenner and Schmulling, 2012). These results suggest that *MYB12* might be part of a negative feedback loop where TMO5/LHW, via increased cytokinin signalling, activates its own repressor to modulate vascular proliferation rates.

**Figure 6.**
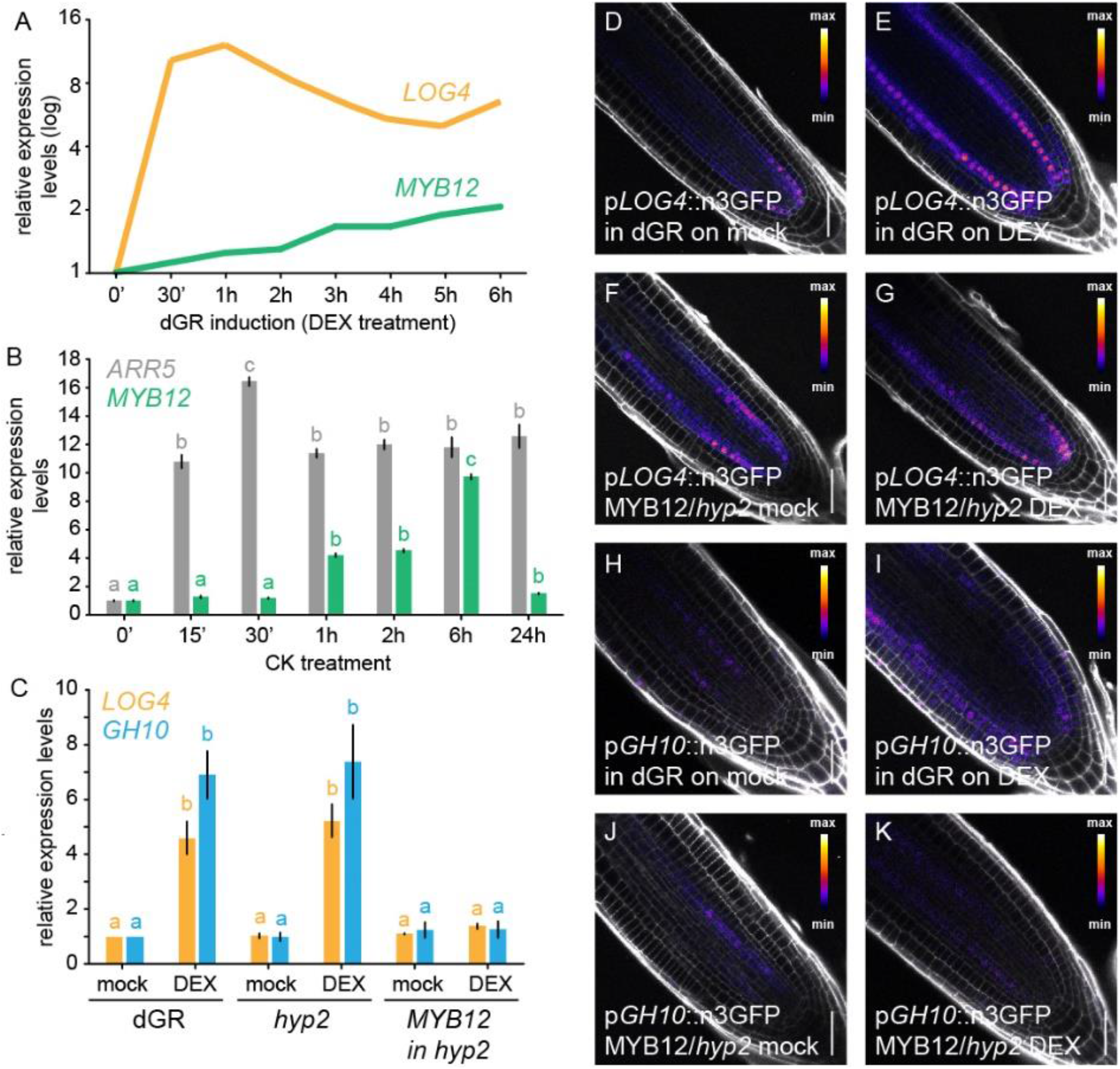
MYB12, downstream of TMO5/LHW-mediated CK production, represses TMO5/LHW target gene expression. (**A**) Relative expression levels *LOG4* and *MYB12* genes over different DEX treatment durations on dGR line derived from microarray data described in Smet et al 2019 (Smet et al., 2019), with 0h DEX expression levels set to 1. (**B**) Relative expression levels of the CK-inducible A-type *ARR5* and *MYB12* in a time course experiment following cytokinin treatment. **(C)** Relative expression of LOG4 and GH10 in 5-days-old seedlings of dGR, hyp2/myb12-2 (dGR) and pRPS5A::MYB12 in hyp2/myb12-2 (dGR) where TMO5/LHW activity was induced for 2h on mock or DEX. (**D-G**) Expression of p*LOG4*::n3GFP in F1 5-days-old seedlings in dGR background (**D**-**E**) and p*RPS5A*::MYB12 in *hyp2/myb12-2* (dGR) (**F**-**G**) background after 24h on mock (**D**,**F**) or DEX (**E**,**G**). Expression of p*GH10*::n3GFP in F1 5-days-old seedlings in dGR background (**H**-**I**) and p*RPS5A*::MYB12 in *hyp2/myb12-2* (dGR) background (**J-K**) after 24h on mock (**H**,**J**) or DEX (**I**,**K**). Scale bars are 50 µm. Lower-case letters in B, C indicate significantly different groups per gene as determined by one-way ANOVA with post-hoc Tukey HSD testing. Black lines indicates mean values and grey boxes indicate data ranges. n marks the number of datapoints for each sample. Error bars are standard errors.

### MYB12 represses TMO5/LHW transcriptional activity

One possible way how MYB12 could repress TMO5/LHW is by altering the downstream cytokinin response. To test this hypothesis, we analysed the inhibition of root length caused by increasing concentrations of exogenously applied cytokinin in *myb12* mutants. No major differences in cytokinin sensitivity were observed between either *myb12* allele and their respective control lines under mock conditions (Col-0 for *myb12-1f* and dGR for *hyp2/myb12-2*) (**Fig. S5, Table S1**), suggesting the repression of TMO5/LHW activity does not act at the level of cytokinin signalling or perception. Next, we tested possible repression at the level of the activity of the TMO5/LHW heterodimer itself by analysing the expression levels of direct TMO5/LHW target genes in the *hyp2/myb12-2* and p*RPS5A*::MYB12 *hyp2/myb12-2* dGR lines in comparison to the dGR control. The expression levels of the direct target genes *LOG4* and *GH10* can be used as molecular read-out of TMO5/LHW activity (De Rybel et al., 2014; Ohashi-Ito et al., 2014; Vera-Sirera et al., 2015). Upon DEX treatment, relative expression levels of *LOG4* and *GH10* were induced in control (dGR) and *hyp2/myb12-2* in dGR backgrounds (**Fig. 6C, Table S1**). In the p*RPS5A*::MYB12 *hyp2/myb12-2* dGR line, however, no induction in *LOG4* and *GH10* expression was observed (**Fig. 6C, Table S1**), suggesting that MYB12 might directly inhibit TMO5/LHW activity. To verify these results, we next introduced the transcriptional reporter of *LOG4* (De Rybel et al., 2014) and a newly generated reporter for *GH10* into the p*RPS5A*::MYB12 *hyp2/myb12-2* dGR line and the parental dGR line as control. Both the p*LOG4*::n3GFP and p*GH10*::n3GFP transcriptional reporters showed a clear induction in expression strength and ectopic expression upon DEX treatment in dGR/+ background compared to a mock DMSO treatment (**Fig. 6D-E, H-I**). This induction was repressed in the p*RPS5A*::MYB12/+ *hyp2/+* dGR/+ background (**Fig. 6F-G, J-K**); confirming the qRT-PCR results (**Fig. 6C, Table S1**). Taken together, these results suggest that MYB12 represses TMO5/LHW activity by inhibiting direct target gene expression. Importantly, MYB12 does not contain a characteristic EAR motif associated with transcriptional repressors (Kagale and Rozwadowski, 2011; Liu et al., 2015) and directly activates transcription of the *CHS* and *FLS* genes (Mehrtens et al., 2005). This shows that MYB12 is thus not a typical transcriptional repressor, but represses TMO5/LHW transcriptional activity in another way.

### MYB12 non-competitively binds to TMO5 in xylem cells

TMO5/LHW activity is known to be repressed by the SACL bHLH proteins, which compete with TMO5 for binding to LHW and thus reduce the amount of active TMO5/LHW dimers (Katayama et al., 2015; Vera-Sirera et al., 2015). Given the well documented interactions between MYB and bHLH TFs (Zhao et al., 2008; Carretero-Paulet et al., 2010; Feller et al., 2011; Cui et al., 2021), we hypothesized that MYB12 function might involve direct binding to the TMO5/LHW complex, similar to the SACL mode-of-action (Katayama et al., 2015; Vera-Sirera et al., 2015). Firstly, if MYB12 were to bind the TMO5/LHW complex, it would need to be present in the same cells. Therefore, we determined the spatiotemporal expression domain of *MYB12* by analysing a newly generated p*MYB12*::nGFP/GUS transcriptional reporter line. Although very faint expression was observed in young xylem cells, *MYB12* was found to be broadly and strongly expressed in most cells from the elongation zone onwards; including xylem cells and in the columella cells (**Fig. 7A-B**). This expression domain fits with predictions from a recently published single cell atlas of the root apical meristem (Wendrich et al., 2020) (**Fig. S4C)**. Given that TMO5 and LHW are expressed in xylem cells (De Rybel et al., 2013) (**Fig. 7C-D**), direct *in planta* binding of MYB12 to TMO5/LHW complex is possible.

**Figure 7.**
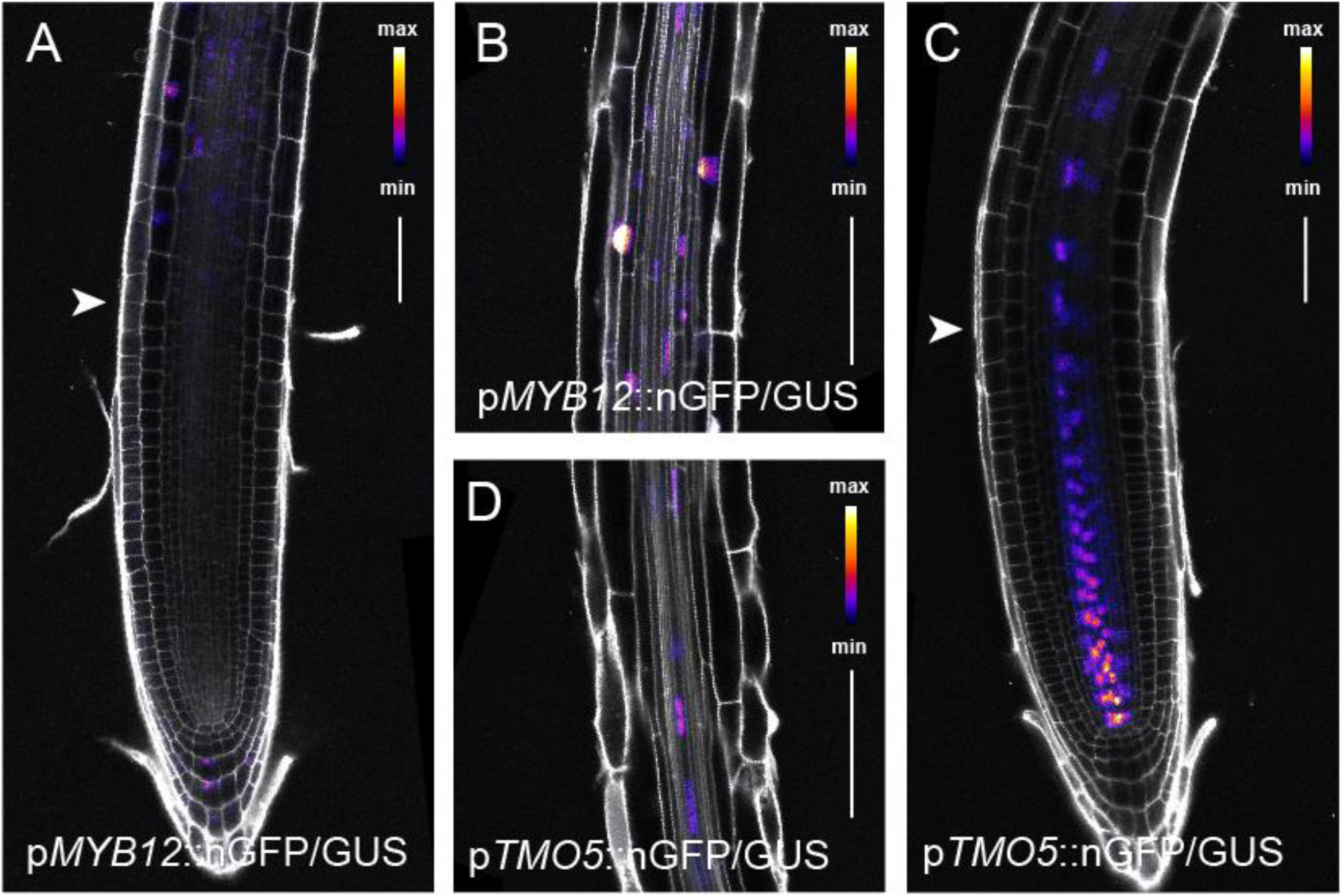
MYB12 and TMO5 have overlapping expression patterns. (**A-B**) Expression pattern of 1-week-old p*MYB12*::nGFP/GUS in root meristem (**A**) and root elongation zone (**B**). (**C**-**D**) Expression pattern of 1-week-old p*TMO5*::nGFP/GUS in meristem(**C**) and root elongation zone (**D**). Arrowheads indicate start of root elongation zone. Scale bars are 50 µm (**A**,**C**) and 100 µm (**B**,**D**).

Next, we thus tested the capacity of MYB12 to directly interact with TMO5 and/or LHW. Yeast-2-Hybrid (Y2H) analysis showed that MYB12 is able to bind to TMO5 (**Fig. 8A**). Binding of MYB12 to LHW could not be evaluated due to auto-activation in the yeast system (**Fig. 8B**). To provide confirmation of this interaction *in planta* using an independent system, we took advantage of the recently developed rapamycin-dependent Knock Sideways assay in transiently transformed *N. benthamiana* leaves (Winkler et al., 2021). This assay is based on the ability of FKBP and FRB protein domains to solely dimerize in presence of the drug rapamycin (Belshaw et al., 1996). In control conditions, we observed that simultaneous infiltration of plasmids carrying TMO5-GFP-FKB, MYB12-TagBFP2 and a mitochondria targeted FRB in *Nicotiana benthamiana* leaves resulted in nuclear localization of the TMO5 and MYB12 fusions, as expected from transcription factors (**Fig. 8C**). In the presence of rapamycin, TMO5-GFP-FKBP bound to mito-FRB and delocalized to the mitochondria; along with MYB12-TagBFP2 (**Fig. 8D**). Taken together, these experiments show that TMO5 and MYB12 can directly interact *in vivo* and *in planta*. We next asked whether the MYB12/TMO5 interaction might prevent the formation of the TMO5/LHW complex; similarly to the competitive binding of LHW by SACL3 and TMO5 (Katayama et al., 2015; Vera-Sirera et al., 2015). To test this hypothesis, we evaluated the interaction between TMO5 and LHW in the presence or absence of MYB12 in Yeast-3-Hybrid (Y3H) experiments (**Fig. 8E**). Although a few colonies occasionally showed auto-activation of TMO5 in presence of MYB12, all tested colonies showed a clear interaction between TMO5 and LHW in presence of MYB12, while the MYB12 negative control showed no growth on selective medium (**Fig. 8E**). We thus failed to identify a competitive inhibition by MYB12 of TMO5/LHW heterodimer formation in our yeast system. Although we have to be careful in concluding from a negative readout, these results suggest that MYB12 might not inhibit TMO5/LHW activity by competitive inhibition of the formation of the TMO5/LHW complex itself, as is the case for the SACL proteins (Katayama et al., 2015; Vera-Sirera et al., 2015).

**Figure 8.**
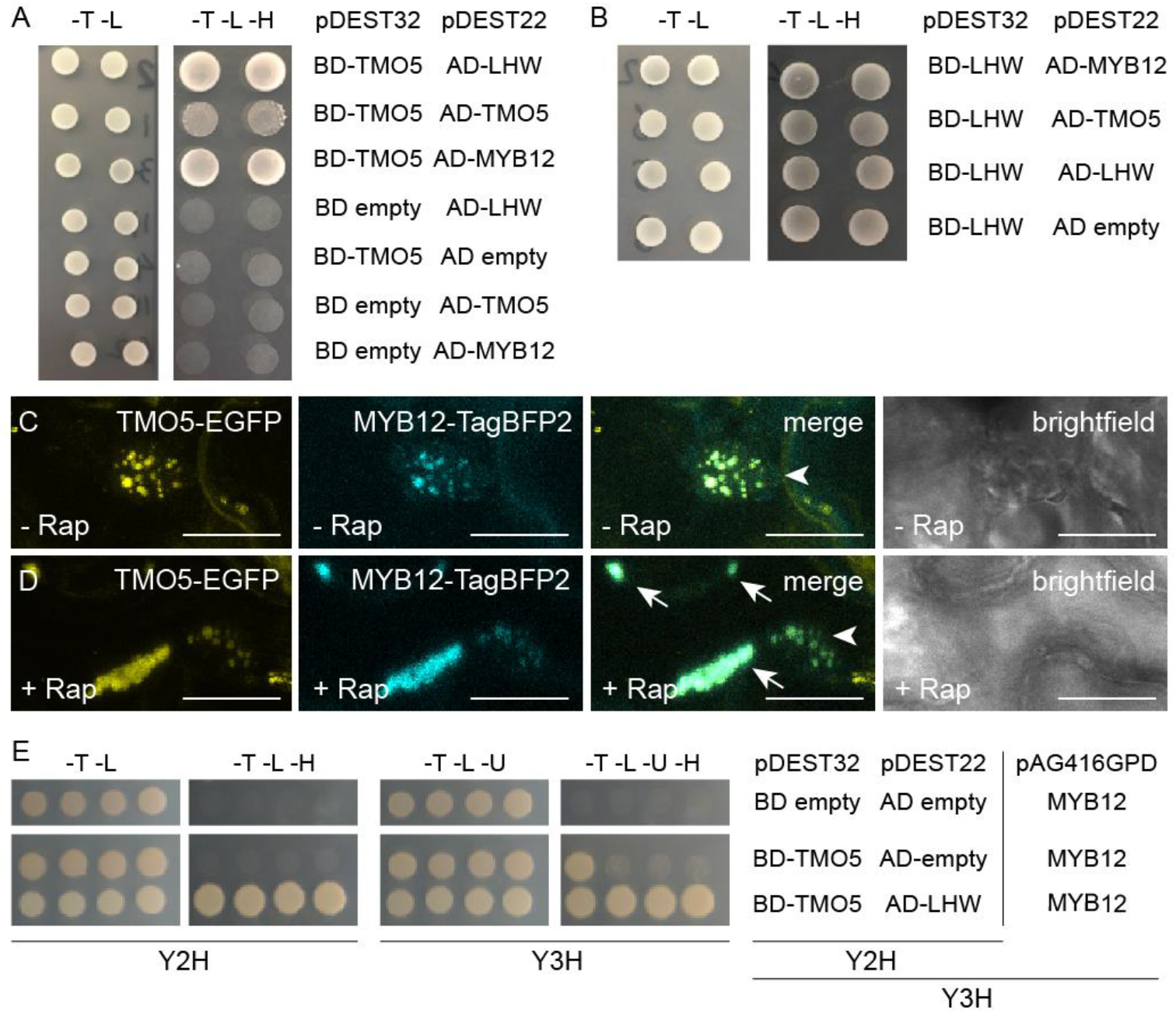
MYB12 binds to TMO5 in yeast and tobacco leaves. (**A-B**) Yeast-two-hybrid assay with pDEST22 (prey) or pDEST32 (bait) constructs containing fusion proteins of the MYB12, TMO5 and LHW coupled to respectively, the activator (AD) or binding domain (BD). The empty pDEST22 or empty pDEST32 plasmids were used to check for auto-activation. Transformed yeast grown on the selective -Trp/-Leu (-T -L) medium and interaction verifying -Trp/-Leu/-His (-T -L -H) medium. (**C-D**) Knock-sideways with TMO5-EGFP-FKGP, MYB12-TagBFP2 and Mito-FRB in absence (**C**) or presence of rapamycin (**D**). Arrows indicate the aggregated mitochondria and arrowheads indicate the nucleus. (n ≥10) Scale bars are 20 µm. (**E**) Left hand side images Y2H yeast pairs of pDEST32 and pDEST22 constructs selective -Trp/-Leu (-T -L) medium and interaction verifying -Trp/-Leu/-His (-T -L - H) medium. MYB12 in pAG416GPD with marker gene URA3 was introduced by mating. Right images Y3H yeast transformants with pDEST32 and pDEST22 constructs and pAG416GPD on selective -Trp/-Leu/-Ura (-T -L -U) medium and interaction verifying selective -Trp/-Leu/-Ura/-His (-T -L -U -H) medium.

Altogether, using forward genetics, we have identified the R2R3 MYB transcription factor MYB12 as a novel regulator of the TMO5/LHW pathway during root vascular proliferation. MYB12 directly interacts with TMO5 and represses the activity of the TMO5/LHW complex at the level of direct target gene expression. *MYB12* itself is a downstream target gene of the TMO5/LHW pathway, thus constituting a negative feedback loop which contributes to fine tuning the activity of the TMO5/LHW complex during vascular development.

## Discussion

The patterning and proliferation of the vascular bundle during primary root growth relies on a complex regulatory network of transcriptional, hormonal and other signals (De Rybel et al., 2016). The key heterodimeric bHLH transcription factor complex, TMO5/LHW, promotes cytokinin biosynthesis through the expression of *LOG3, LOG4* and *BGLU44* in the xylem cells (De Rybel et al., 2014; Ohashi-Ito et al., 2014; Yang et al., 2021). This locally produced cytokinin is thought to act as a mobile signal that coordinates the radial growth and correct patterning of the vascular bundle (Wybouw and De Rybel, 2019). In this study, we have taken a forward genetic approach to find new regulators of the TMO5/LHW pathway and discovered a novel function of the previously described transcription factor MYB12. Our data revealed that *myb12* mutants are hypersensitive to the gain-of-function phenotypes caused by TMO5/LHW misexpression, while *MYB12* misexpression represses vascular proliferation by inhibiting the transcriptional activation of direct TMO5/LHW targets genes. Moreover, *MYB12* is transcriptionally activated by the cytokinin response downstream of TMO5/LHW, and MYB12 directly interacts with TMO5. All these findings indicate that MYB12 acts as a repressor of the *TMO5/LHW* transcriptional pathway, while at the same time being its downstream target. Hence, we have found a novel negative feedback loop regulating the TMO5/LHW transcriptional network via the action of MYB12.

This negative feedback loop is reminiscent of the previously described regulation of the TMO5/LHW pathway by the *SACL* genes. Nonetheless, there are several key differences between the MYB12-and SACL-mediated negative feedback loops. Firstly, *MYB12* appears to be a secondary TMO5/LHW target induced indirectly by the downstream cytokinin response, while the *SACL* genes are direct targets of TMO5/LHW (Katayama et al., 2015; Vera-Sirera et al., 2015). This would suggest that the MYB12-mediated negative feedback is slower in comparison to the SACL loop, which might be important for spatiotemporal fine tuning TMO5/LHW activity. Furthermore, the cytokinin response levels are affected by numerous factors other than TMO5/LHW (Kieber and Schaller, 2018). Thus, the cytokinin-inducible MYB12 can, unlike the SACL proteins, help optimize vascular proliferation rates by integrating the TMO5/LHW activity with other developmental signals. In support of the SACL-and MYB12-mediated negative feedback loops acting on different spatiotemporal scales, *SACL* and *MYB12* have very distinct expression patterns. *SACL*s are co-expressed with *TMO5* and *LHW* in xylem cells in the root meristem zone (Vera-Sirera et al., 2015). *MYB12* is most prominently expressed in older root tissues from the differentiation zone onwards, consistent with providing slower and more indirect feedback. However, the SACL and MYB12 regulatory loops do not seem to be mutual exclusive, as *myb12* mutants are hypersensitive towards increased TMO5/LHW activity in the root meristem. Unfortunately, despite clear inhibitory effects on vascular proliferation in both *SACL* and *MYB12* gain-of-function lines, a lack of prominent aberrant phenotypes in the respective loss-of-function mutants makes it difficult to dissect the exact function of these genes during vascular development. This further emphasizes the pronounced genetic redundancy operating in plant development, especially during the control of such vital processes like vascular tissue patterning.

We have shown that MYB12 directly interacts with TMO5 and inhibits the transcriptional activation of direct TMO5/LHW target genes. Nonetheless, the exact molecular mechanism of MYB12 action remains partially unclear. In yeast, we could not show that MYB12 disrupts the TMO5/LHW complex formation like the SACLs do (Katayama et al., 2015; Vera-Sirera et al., 2015). Moreover, it does not contain an EAR or TLLLFR motif typical for MYB TF repressors (Kagale and Rozwadowski, 2011; Ma and Constabel, 2019). Additionally, MYB12 lacks the bHLH-binding motif present in other known bHLH-interacting MYB TFs (Zimmermann et al., 2004; Wang and Chen, 2014), and it functions as a bona fide transcriptional activator in other developmental contexts (Forkmann and Martens, 2001; Mehrtens et al., 2005). Therefore, the MYB12-mediated inhibition of TMO5/LHW activity must depend on another molecular mechanism.

In one scenario, MYB12 might act as a passive repressor by preventing TMO5/LHW interaction with DNA and/or recruitment of the RNA polymerase II complex (Kazan, 2006; Krogan and Long, 2009). Another and more likely possibility is that rather than acting as a conventional repressor, MYB12 might redirect TMO5/LHW activity away from *LOG4, GH10* and other genes involved in vascular proliferation, and contribute to activating different TMO5/LHW target genes instead. This explanation would fit best with the previously described function of MYB12 as a classical transcriptional activator of several genes in the flavonoid biosynthesis pathway (Forkmann and Martens, 2001; Mehrtens et al., 2005). Target gene specificity has previously been associated with the MYB TFs in heteromeric bHLH-MYB transcriptional complexes (Ramsay and Glover, 2005). TMO5-LIKE 1 (T5L1), a close homolog of TMO5, is able to promote ectopic xylem differentiation in addition to its role in promoting radial growth (Katayama et al., 2015); The same bHLH TF thus functions in two very different developmental processes that require the activation of completely different gene sets. It is conceivable that such alternative functionalities of bHLH TFs could be achieved by interactions with different MYBs. In such a scenario, the TMO5/LHW complex would recruit an unknown MYB TF to promote the expression of genes required for vascular proliferation, while the alternative recruitment of MYB12 would lead to the activation of different target genes. To take this speculation even further, the dual roles of MYB12 in flavonol biosynthesis (Forkmann and Martens, 2001; Mehrtens et al., 2005) and vascular proliferation (this study) could then be explained by alternative interactions with TMO5 and an unknown bHLH TF needed for MYB12-mediated induction of the *CHS* and *FLS* flavonol biosynthesis genes. Further investigations into the precise molecular mechanisms responsible for MYB12 as well as other related MYB TFs will be needed to shed light on these intriguing open questions and hypotheses.

What is the biological meaning of the same transcription factor MYB12 being involved in flavonol biosynthesis as well as vascular proliferation is another open question arising from our study. Interestingly, the bHLH TF TRANSPARENT TESTA 8 (TT8) has been previously implied in flavonoid biosynthesis (Nesi et al., 2000) and trichome development (Maes et al., 2008), indicating that dual functions in different metabolic and developmental pathways might be a common feature of multiple transcription factors from different families. This might reflect the need of certain metabolic changes for a specific developmental process. For example, trichomes are rich in biotic stress defence compounds which include flavonoids (Karabourniotis et al., 2020). Utilizing TT8 to control both trichome development and flavonoid biosynthesis might thus aid in coordinating the two processes. Likewise, the transition from vascular proliferation to differentiation might involve so far unappreciated metabolic changes in addition to the decline of TMO5/LHW activity, both hypothetically controlled by MYB12. Alternatively, dampening the TMO5/LHW pathway while promoting flavonoid biosynthesis might contribute to the balance between growth and defence processes. Different stresses often lead to increased reactive oxygen species levels, which can be mitigated by flavonoid antioxidant activity (Wang et al., 2016). In such conditions, attenuating the TMO5/LHW-mediated radial growth in favour of flavonoid biosynthesis by the increased MYB12 levels could be important for optimal resource allocation.

In summary, we have uncovered a novel role of the transcription factor MYB12 as a negative regulator of the TMO5/LHW pathway during vascular proliferation. The MYB12-mediated negative feedback loop is distinct from the modus operandi of the previously described SACL proteins in both molecular mechanism and spatiotemporal dynamics, showing that TMO5/LHW activity is being controlled using multiple distinct mechanisms. The full molecular details of MYB12 mode of action, as well as the biological meaning of its dual functions in vascular development and flavonoid biosynthesis, remain exciting challenges for future investigations. Our work establishes that a bona fide transcriptional activator can function as a repressor in a different transcriptional network. Furthermore, our results show that functional interactions between bHLH and MYB transcription factors are involved in multiple unrelated transcriptional networks, highlighting them as a powerful and possibly underappreciated developmental module.

## Materials and Methods

### Plant material and growth conditions

Seedlings were grown at 22°C under continuous light on ½ Murashige and Skoog (MS) medium without sucrose, after seeds were stratified for 24h-48h. For dexamethasone (DEX) treatment, 10 µM DEX (Sigma-Aldrich) was added to the growth medium from a 10 mM DMSO stock solution; seedlings were either germinated on DEX-containing medium or transferred from MS medium at the indicated time point. For the CK sensitivity assay, seedlings were germinated on 6-benzylaminopurine (6-BAP; Duchefa) -containing medium. The AGI identifiers for the genes used in this manuscript are as followed: *TMO5 (AT3G25710), LHW (AT2G27230), MYB12* (*AT2G47460*), *LOG4* (*AT3G53450*), *GH10* (*AT4G38650*) and *ARR5* (*AT3G48100*). The following mutant and transgenic lines were described previously: *myb12-1f* (Mehrtens et al., 2005); *myb11 myb12-1f myb111* (*myb* triple) (Stracke et al., 2007); p*RPS5A*::TMO5:GR x p*RPS5A*::LHW:GR (dGR) (Smet et al., 2019); p*LOG4*::n3GFP (De Rybel et al., 2014). The lines *ins4/lhw-8* and *hyp2/myb12-2* were generated in the dGR background by EMS mutagenesis (see below). The lines p*GH10*::n3GFP, p*RPS5A*::MYB12, p*RPS5A*::MYB12 *hyp2* and p*MYB12*::nGFP-GUS were obtained by transforming the respective expression clones into Col-0 or *hyp2* by the floral dip method (Clough and Bent, 1998). The p*LOG4*::n3GFP and p*GH10*::n3GFP were introduced into the dGR and p*RPS5A*::MYB12 *hyp2/myb12-2* dGR backgrounds by genetic crossing and analysed in F1 generation seedlings.

### EMS mutagenesis and screening

The dGR line (Smet et al., 2019) was used for the EMS mutagenesis. Approximately 10,000 seeds were incubated shaking in water overnight. The water was replaced with 15 ml of 0.05 % Triton X-100. After mixing well, the seeds were incubated for 5 min in this solution then twice washed with water. The seeds were mutagenized by treatment with 30 mM EMS in 0.1 M phosphate buffer (pH 7.5) for 6-7 hours. Afterwards, the EMS solution was removed, and mutagenesis was stopped by adding 0.1 M Na_2_S_2_O_3_ for 5 min five times. The Na_2_S_2_O_3_ was washed away with water seven times. These seeds were afterwards stratified in 0.1% agarose overnight. Approximately 50 seeds were sown together in a pot per pool. A total of 228 pools was maintained. For each pool, about 1,000 M_2_ seeds were initially screened on 10 µM DEX containing ½ MS media, leading to a selection of 260 mutants from 110 pools. Next, the root length and root width of one-week-old M_3_ seedlings was measured in both mock (DMSO) and 10 µM DEX. Changes in root length and meristem width were measured upon DEX treatment and compared to a Col-0 and dGR control.

### Mapping causal mutation of EMS mutants

Selected EMS mutants were backcrossed with the parental dGR line, and one-week-old BC_1_F_2_ seedlings with the desired phenotype were collected for DNA extraction. DNA was extracted using hexadecyltrimethylammoniumbromide (CTAB) extraction buffer (0.1 M Tris pH7.5, 0.7M NaCl, 0.01 M EDTA and 0.03 M CTAB) and afterwards isolated using chloroform:isoamylalcohol (24:1) and isopropanol. RNA was degraded by RNase treatment between the chloroform and isopropanol isolation steps. The bulked genomic DNA was sequenced by using the Illumina NextSeq 500 system. For the library preparation, an insert size of 400-500 bp was used. Paired end sequencing was performed, with a read length of 2×150 bp length and 50x coverage. Potential causal mutations are selected by using the SHORE map analysis tool (Schneeberger et al., 2009).

### Molecular cloning

The promotors and coding sequences were PCR amplified using a high-fidelity polymerase (primers used are shown in **Table S3**). All constructs were made by MultiSite Gateway cloning (Karimi et al., 2002). Promoter regions were amplified from genomic DNA and introduced into the *pDONRP4P1R* vector. The coding sequences were amplified from root cDNA and introduced into the *pDONR221* vector. All entry clones were sequence verified before further steps. The MYB12 promoter entry was cloned into pmK7S*nF14mGW destination vector. The construct was transformed in Col-0 and dGR via Agrobacterium mediated flower dipping (Clough and Bent, 1998).

### Root phenotyping

For root length measurements, one-week-old roots were scanned on a flatbed scanner and root length was measured by using the freeware program FIJI with the integrated NEURONJ plugin (https://imagescience.org/meijering/software/neuronj/) (Meijering et al., 2004). Root width of one-week-old seedlings were measured by dissecting the roots and mounting them in clearing agent (60 % lactic acid, 20 % glycerol and 20 % H_2_O). Width of the root tips was measured at the beginning of the elongation zone for all roots by using FIJI (Schindelin et al., 2012). Imaging of differentiated primary xylem vessels was performed on one-week-old roots mounted in the clearing agent described above.

### Statistics and visualization of the data

All boxplots were generated with BoxPlotR web tool (http://shiny.chemgrid.org/boxplotr). In these plots, the boxes indicate the median, 25th and 75th percentile of the data, the whiskers extend to minima and maxima within 2 SDs of the mean, and outliers are indicated as single empty circles. ‘n’ represents the number of data points. Pairwise comparisons were performed using standard two-sided Student’s T-testing. Student T-test significances asterisks: * = p-value < 0.05; ** = p-value < 0.01; *** = p-value < 0.001. The lower-case letters associated with the boxplots indicate significantly different groups as determined by one-way or two-way ANOVA with post-hoc Tukey HSD testing (p<0.05).

### Confocal imaging

Transcriptional and translational fluorescent reporter lines were imaged on a Leica SP8 confocal microscope with a 40x NA 1.1 water immersion objective. Seedlings were mounted in propidium iodide (PI); GFP and sYFP reporter lines were excited at 488, resp. 514 nm and detected at 500-535, resp. 515-550 nm; PI was detected at 600-700 nm. For the vascular cell file number measurements, one-week-old seedlings were fixed and stained using the mPS-PI protocol and imaged using the Leica SP2 or SP8 confocal microscopes as described previously (Truernit et al., 2008; Arents et al., 2022). The vascular bundle cell number quantifications included the pericycle cell layer, except if mentioned otherwise.

### RNA isolation and qRT-PCR

For dGR induction, plants were grown on ½ MS (1% agar) for 5 days before transferring to either mock or 10 µM DEX for 2h. For CK treatment, 5-day-old seedlings were transferred to medium containing 10 µM 6-benzylaminopurine (6-BAP; Duchefa) from a 10 mM DMSO stock solution. All samples were ground in liquid nitrogen and RNA was extracted using RNA isolation protocol for non-fiberous tissue by the RNA Tissue Miniprep System (Promega). cDNA synthesis was done using 1µg of total RNA with the qScriptTM cDNA Supermix kit (Quanta BioSciences). The qRT-PCR primers were designed by Universal Probe Library Design Center (Roche) (**Table S3**). The qRT-PCR was performed using *UBC* and *EEF* as reference genes on a Roche Lightcycler 480 device (Roche Molecular Systems Inc.) with SYBR Green I Master kit (Roche). The gene expression analysis was done using qBase v3.2 software (Biogazelle, Zwijnaarde, Belgium - www.qbaseplus.com).

### DNA extraction and genotyping

Genomic DNA was isolated using the CTAB extraction method. The T-DNA mutants (*myb11*/SALK077068 and *myb111*/GK291D01) were genotyped using PCR based method (**Table S3**). The *myb12-1f* mutant (Mehrtens et al., 2005) was genotyped using cleaved amplified polymorphic sequence (CAPS). An amplicon of 547 bp was amplified (using primers described in **Table S3**), and was cut by using HphI restriction. The wild type allele is cut into two bands of 399 bp and 148 bp, while the mutant remained uncut.

### Yeast-2-Hybrid (Y2H) and Yeast-3-Hybrid (Y3H) analysis

The MYB12, TMO5 and LHW coding sequences were cloned into pDEST22 (Prey: GAL4AD-x Yeast selection marker: TRP1) and pDEST32 (Bait: GAL4DB-y Yeast selection marker: LEU2). These plasmids were transformed into *Saccharomyces cerevisiae* strain AH109 (Clontech). At least 3 independent yeast transformants were checked for each pairwise interaction according to (Cuellar et al., 2013) with minor modifications: the protein-protein interactions were validated with undiluted overnight yeast culture droplets manually pipetted on selective SD Base-Leu/-Trp/-His and grown for 3-4 days at 30°C before imaging. The Y3H was performed as described in Yperman *et al* (2021) (Yperman et al., 2021). For the Y3H, bait and prey subunits (TMO5 and LHW) were cloned in pDEST32 and pDEST22 expression vectors and transformed via heat shock in the PJ69-4α yeast strain. MYB12 was cloned in pAG416GPD (Yeast selection marker: URA3) and transformed via heat shock in the PJ69-4a yeast strain. A and α strains were mated and cultured in SD–Leu/−Trp/–Ura. Cultures were grown for 2 days and were diluted to OD600 0.2, and 10 µl was pipetted on SD–Leu/−Trp/–Ura and SD–Leu/−Trp/–Ura/-His and grown for 3 days at 30°C, after which the plates were imaged.

### Knock sideways

The knock sideways (KSD) assay was performed as described previously [52]. Briefly, *N. benthamiana* leaves were transiently transformed with the constructs p*G1090::XVE>>*MYB12-TagBFP2, p*35S*::TMO5-EGFP-FKBP and p*35S*::MITO-FRB. After ca. 24h, the transformed leaves were infiltrated with 1 µM rapamycin or H_2_O mock control. Images were acquired 24-30 h thereafter on a Leica SP8X confocal microscope in line sequential scanning mode. The p*G1090::XVE>>*MYB12-TagBFP2 was originally intended for estradiol-inducible expression, but turned out very leaky in expression in the *N. benthamiana* system and was thus used for constitutive expression instead.

## Acknowledgements

The authors thank Ralf Stracke and Sofie Goormachtig for sharing the *myb111myb12-1f,myb111* and *myb12-f* mutant seeds, respectively; Davy Opdenacker for help with the EMS mutagenesis and Frederik Coppens for help with the SHOREmap analysis. This work was funded by The Research Foundation -Flanders (FWO; Odysseus II G0D0515N and post-doc fellowship 1215820N); the Netherlands Organization for Scientific Research (NWO; VIDI 864.13.00); the European Research Council (ERC Starting Grant TORPEDO; 714055); the European Union’s Horizon 2020 research and innovation programme under the Marie Sklodowska-Curie grant agreement No 885979 “DIVISION BELL”; EMBO (long term fellowship ALTF 1005-2019); and ERC consolidator Grant T-REX; 682436 to DVD.

## Author Contributions

B.D.R. conceived the project; B.D.R., M.G., D.V.D., B.W. and H.E.A. designed experiments; B.W. and W.S. performed EMS mutagenesis; B.W. and H.E.A. performed the EMS screening; B.W. performed the SHOREmap analysis; B.W. and H.E.A. analysed the role of MYB12; B.Y. and J.N. performed the Y2H experiments; M.G. and J.N. performed the knock sideways experiments; M.V. performed the Y3H experiments; B.D.R. and M.G. supervised the project and wrote the paper with input of all authors.

---

**Figure S1.**
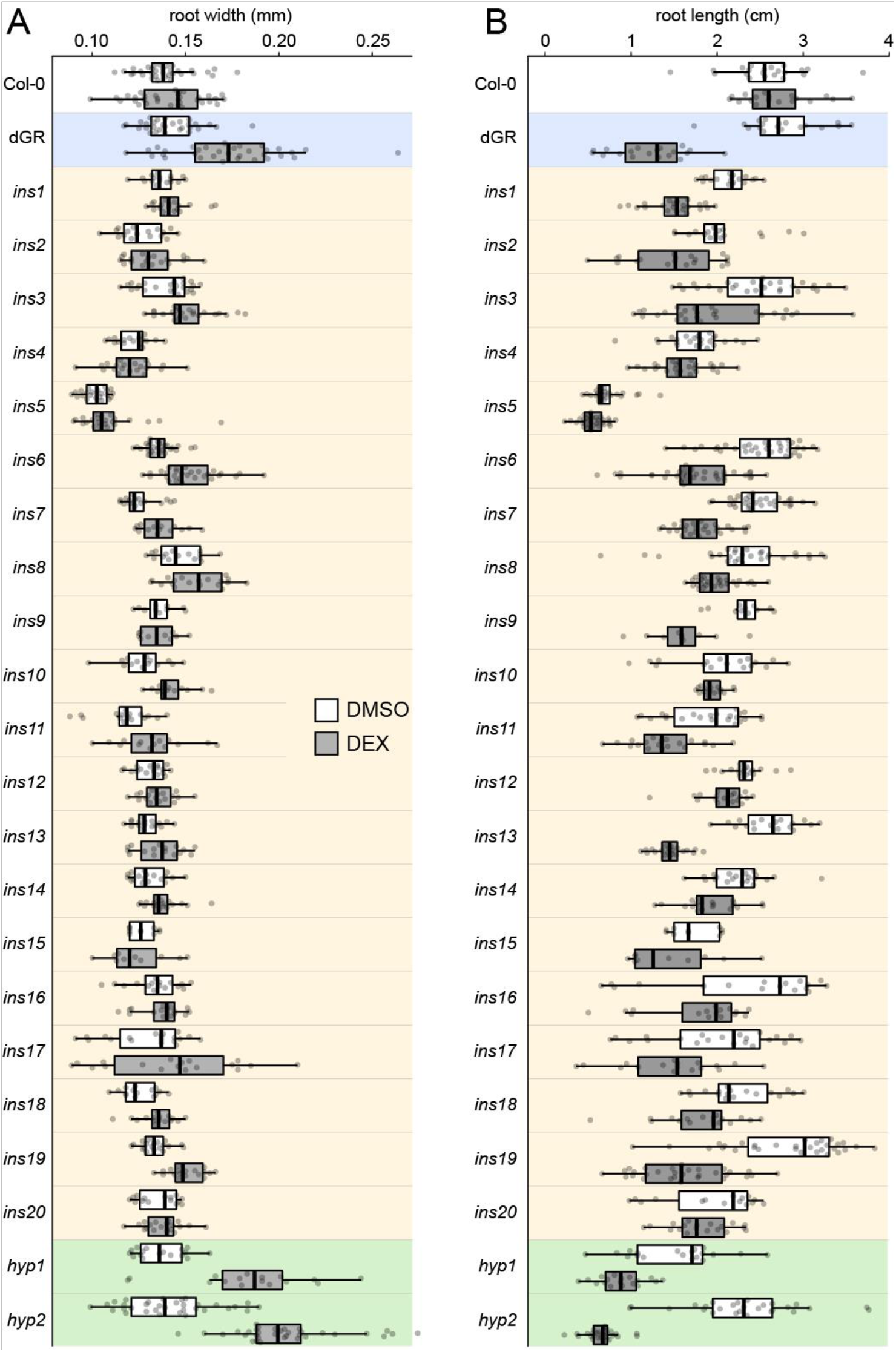
Root widths and lengths of selected EMS mutants. (**A**-**B**) Mutants seedlings were grown on mock or 10 μM DEX for 1-week, were analysed for width (**A**) and length (**B**). Samples were compared pairwise in a two-way ANOVA and post hoc comparison with results shown in **Table S1**.

**Figure S2.**
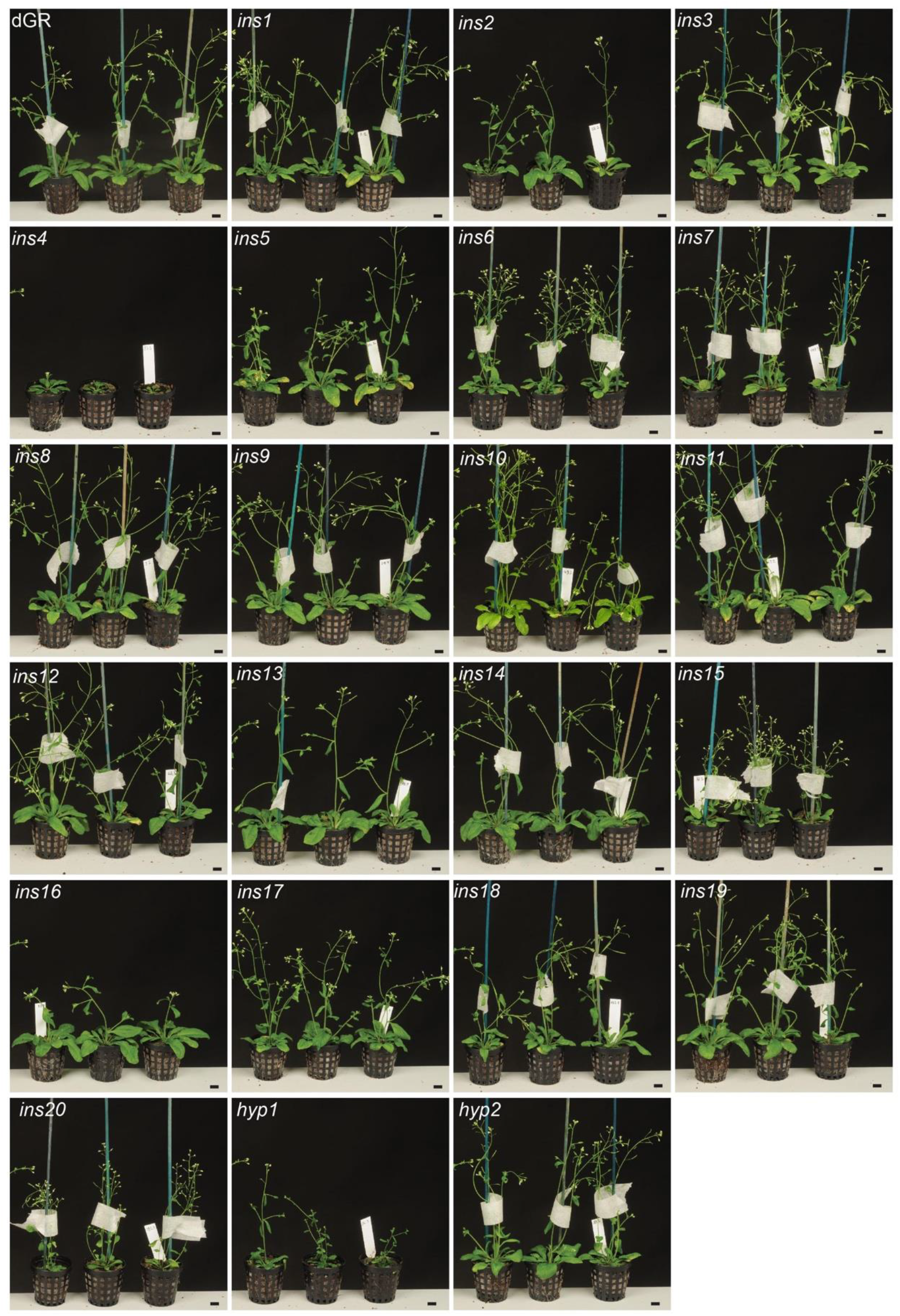
Whole plant phenotype of selected mutants. Overview of selected EMS mutants at 5-week-old. Scale bars are 1 cm.

**Figure S3.**
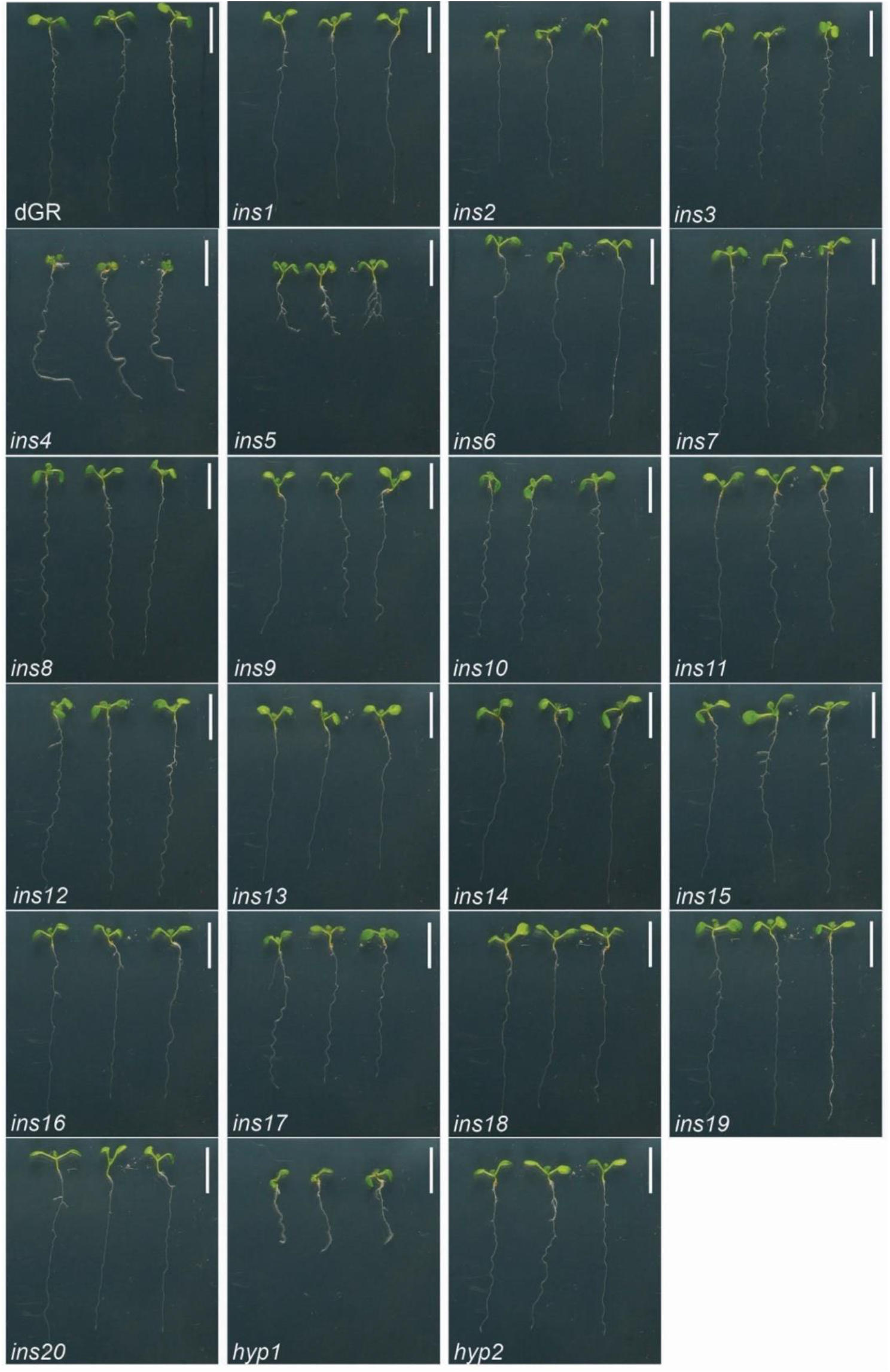
Whole seedling phenotype of selected mutants. Overview of selected EMS mutants at the 1-week-old seedling stage grown on ½ MS medium. Scale bars are 1 cm.

**Figure S4.**
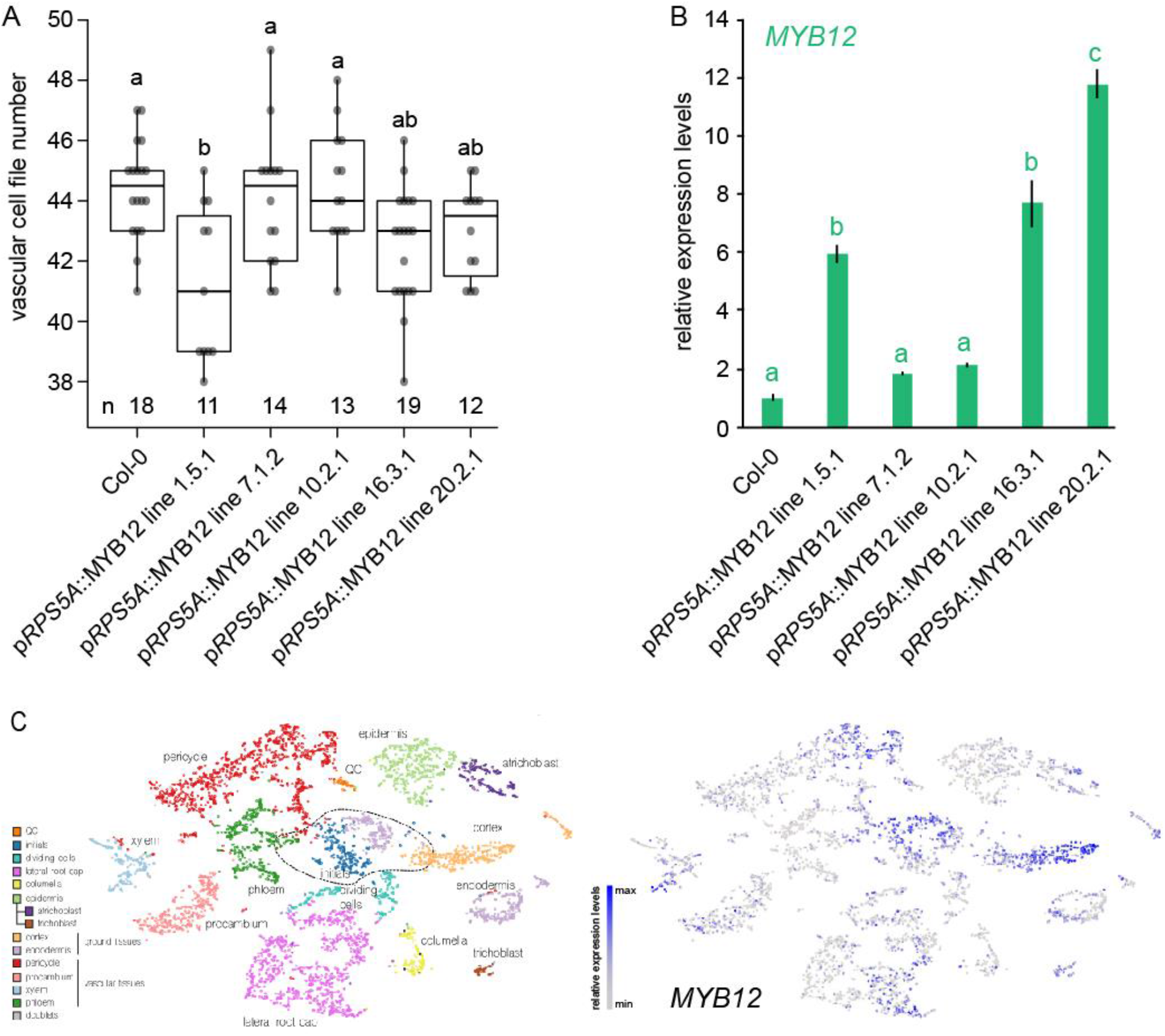
Vascular cell file number in p*RPS5A*::MYB lines and *MYB12* expression levels. (**A**) Vascular cell number in the root meristem of 1-week-old p*RPS5A*::MYB12 in Col-0 seedlings grown on ½ MS. (**B**) Relative expression levels of *MYB12* in p*RPS5A*::MYB12 lines. Lower-case letters in A and B indicate significantly different groups as determined by one-way ANOVA with post-hoc Tukey HSD testing and Tukey-Kramer Grouping. Black lines indicates mean values and white boxes indicate data ranges. n marks the number of datapoints for each sample. (**C**) Predicted expression of *MYB12* according to a previously published single cell atlas of the Arabidopsis root meristem (Wendrich et al., 2020).

**Figure S5.**
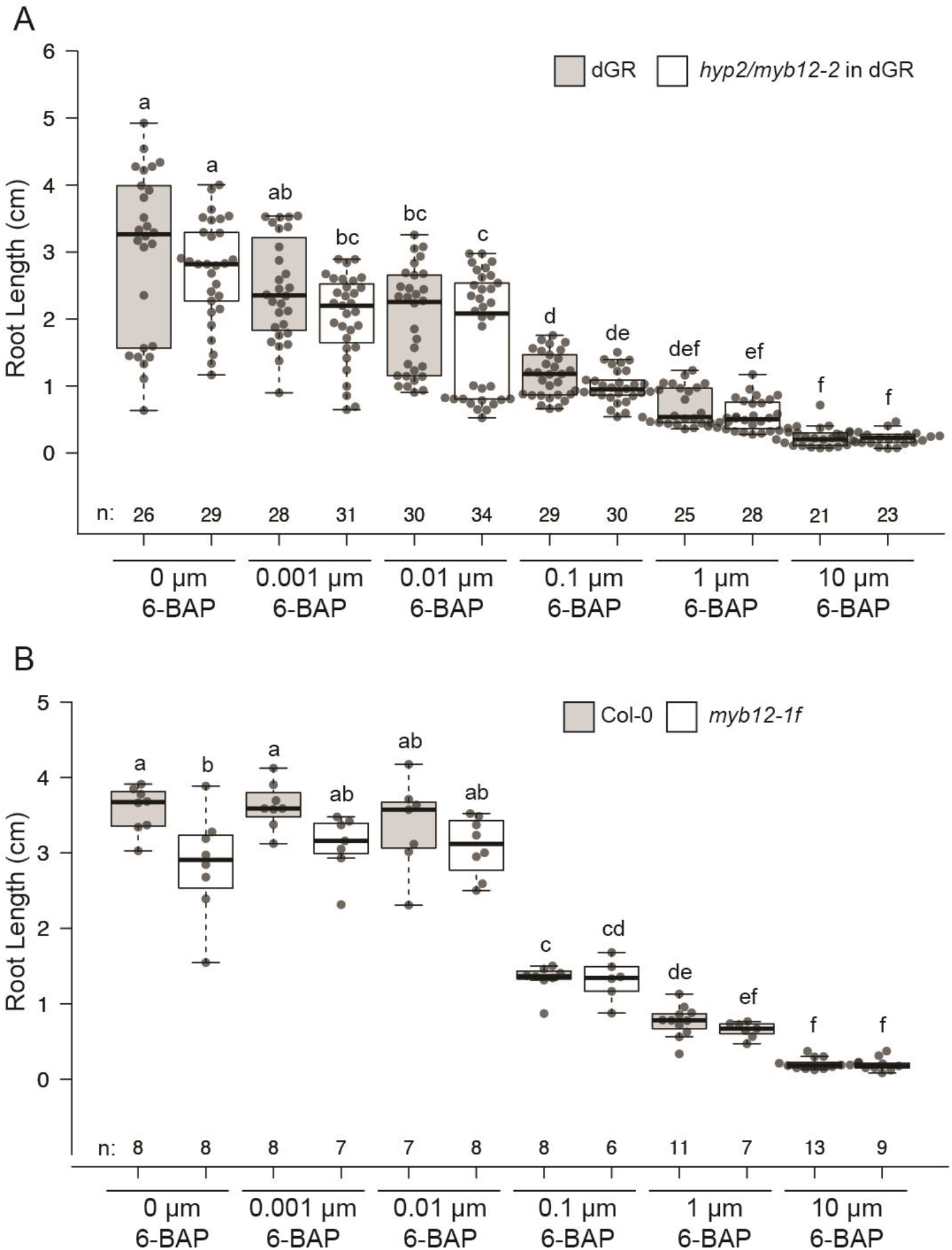
Influence of increasing concentrations of cytokinin on root length. (**A-B**) Root length of 1-week-old seedlings of the indicated genotype grown on ½ MS supplemented with the indicated concentration of cytokinin (6-BAP) from germination onwards. dGR and Col-0 act as controls in panes A and B, respectively. Lower-case letters in A and B indicate significantly different groups as determined by one-way ANOVA with post-hoc Tukey HSD testing and Tukey-Kramer Grouping. Black lines indicates mean values and white/grey boxes indicate data ranges. n marks the number of datapoints for each sample.

